# Spatiotemporal Characterization of Amyloidosis-Associated Microglial States Reveals Sex Difference in Early Plaque Formation

**DOI:** 10.64898/2026.04.05.716600

**Authors:** Amritha Vinayak Manjally, Abigail M. Fowler, Jiratchaya Thanayangyuen, Maximillian Cheval, Monica Iordanov, Daisy Liljegren, Yevah Milord, Jonathan Park, Ema Yamashita, Aidan Conley Kieffer, Tuan Leng Tay

## Abstract

Twice as many women develop Alzheimer’s disease (AD) compared to men. Several key aspects, such as genetic risk factors, hormonal vulnerability, social responsibilities, and differences in longevity, contribute to the strong female bias in AD. To assess whether sex differences can be detected during the onset of AD, we examined the amyloid-β (Aβ) plaque burden––one of the hallmarks of AD––and microglial states in young 5XFAD mouse models of amyloid pathology. We hypothesized that an elevated Aβ burden will directly correlate with an increase in microglial cell number and phagocytic activity, shaping the appearance of compact dense-core plaques in the cortex from 2 to 6 months of age. As expected, no change in microglial density and phenotype was found in Aβ plaque-free hypothalamus of 5XFAD male and female mice when compared to age-matched wildtype controls. By quantifying the number and coverage of diffuse and dense-core plaques in the cortex, we discovered a pronounced increase in Aβ plaques and microglial clustering in 4-month-old female 5XFAD compared to male mice. Conversely by 6 months, higher total plaque load in males and no sex difference in the number of plaque-associated microglial (PAM) cluster was observed. Our spatiotemporal characterization of microglial CD68, Dectin-1 (*Clec7a*), and CD11c (*Itgax*) expression revealed transient sex differences in the upregulation of only CD68 among these phagocytic markers in cortical microglia. In 4-months-old males, greater phagolysosomal activity may benefit Aβ clearance and delay plaque formation. In females, lower microglial phagolysosomal activity and increased microgliosis may have led to the increase in plaque deposition and compaction from 2-4 months. Our results suggest that during early amyloidosis, sex differences in CD68-associated phagolysosomal activity and microglia-driven plaque compaction may cause disproportionate AD risk and severity that is compounded by other exacerbating factors during aging. Taken together, sex-specific targeting of microglial proliferation and phagocytic activity may be a promising intervention in presymptomatic patients with known AD risks.

## INTRODUCTION

By 2025, approximately 4.36 million women and 2.81 million men older than 65 years of age in the United States are estimated to have been diagnosed with clinical AD (Rajan et al., 2021). Women experience faster cognitive decline than men, reportedly with gender specific differences in cognitive profiles in the range of mild to moderate AD (Filon et al., 2016; Holland et al., 2013; Pusswald et al., 2015). One of the major genetic risk factors is *APOE4*, which puts women at a higher risk of developing AD than men (Altmann et al., 2014; Frigerio et al., 2019; Hohman et al., 2018). Estrogen regulates many AD-related genes––including *APOE*––and genomic analyses indicate that menopausal loss of estrogen could increase women’s vulnerability to AD (Ratnakumar et al., 2019).

Progressive extracellular accumulation of Aβ peptides to form plaques is a defining pathological hallmark of AD that contributes to downstream processes including synaptic dysfunction, neuroinflammation, and tau pathology (Lee et al., 2024; McCarter et al., 2013). Diffuse plaques consist of loosely organized, predominantly prefibrillar Aβ aggregates that lack a compact amyloid core. These plaques are relatively common in cognitively unimpaired individuals with amyloid positivity, suggesting that they may represent an early stage of amyloid plaque development (Selkoe and Hardy, 2016).

Increasing evidence demonstrate that microglia play a critical role in the transition from diffuse to dense-core plaque morphology (Baik et al., 2016; Jung et al., 2015). Microglia respond dynamically to Aβ from the earliest stages of AD pathogenesis. Transcriptional and functional studies have identified distinct, stage-specific microglial states, such as disease-associated microglia (DAM), that are driven by the TREM2–APOE signaling pathway and related to phagocytosis-linked genes including *Clec7a* and *Itgax* (Keren-Shaul et al., 2017; Krasemann et al., 2017). Pharmacological and genetic manipulations of microglia in depletion-repopulation models have demonstrated that microglia shape plaque compaction and distribution. Notably, these effects depend on the timing of intervention relative to Aβ plaque emergence (Condello et al., 2015; Spangenberg et al., 2019). A study of Aβ plaque pathology and microglia-plaque interaction within the entorhinal cortex of both sexes revealed greater plaque volume and lower plaque compaction in female 5XFAD AD mice. In addition, the corresponding increase in DAM phenotype and microglial interferon response in female mice was reportedly independent of the estrous cycle (Calcines-Rodríguez et al., 2025). Given that sexual dimorphism is a key factor that underlies microglial development and physiology in the healthy brain (Guneykaya et al., 2018; Hanamsagar et al., 2017; Thion et al., 2018; Villa et al., 2018), we investigate the existence of sex difference in microglial response to the onset of extracellular Aβ deposition and their role in the formation of dense-core plaques.

In this study, we examined the progression of amyloid deposition and compaction in tandem with the changes in DAM and expression of phagocytic markers CD68, Dectin-1 (*Clec7a*), and CD11c (*Itgax*) in 5XFAD male and female mice from 2 to 6 months of age compared to wildtype (WT) controls. To distinguish microglia-plaque interaction during Aβ plaque formation, we performed a unique combinatorial analysis of 6E10 immunostaining for Aβ peptides and compact plaques and Thioflavin S (ThioS) that stains Aβ sheets (Bussière et al., 2004; Shin et al., 2021). Diffuse plaques were identified by 6E10 and the absence of compact ThioS signals while dense-core plaques were detected by both 6E10 and ThioS staining. As expected, we observed that cortical amyloid pathology was nearly absent at 2 months in both sexes of 5XFAD mice. However, the number of both diffuse and dense-core plaques as well as clusters of plaque-associated microglia (PAM) was remarkably increased in females by 4 months. We found higher microglial CD68 expression in non-PAM 5XFAD males than females at 4 months, indicating more efficient Aβ clearance before plaque compaction. Interestingly, no sex difference was observed in in cortical microglial Dectin-1 and CD11c expression at the stages of analysis. By 6 months, higher total plaque burden was observed in males. During early amyloid pathogenesis in 5XFAD mice, microglial upregulation of major histocompatibility complex class II (MHCII) remained low in both sexes. Our findings of early sex-specific phagolysosomal activity in amyloidosis-vulnerable brain regions suggest that temporal sex difference in plaque burden depends on the kinetics of microglial proliferation, formation of PAM and regulation of phagocytosis. Altogether, this reveals an opportunity to target microglial activity as a personalized preventative intervention to delay the onset of amyloid pathology in presymptomatic adults with high risks for AD.

## MATERIALS AND METHODS

### Animals and sample preparation

Wild type C57BL6 and 5XFAD (Oakley et al., 2006) mice were obtained from The Jackson Laboratory (USA) and maintained at the Boston University (BU) Animal Science Center. The mouse strain used for this research project, B6J.SJL-Tg(APPSwFlLon,PSEN1*M146L*L286V) 6799Vas/Mmjax, RRID:MMRRC_034848-JAX, was obtained from the Mutant Mouse Resource and Research Center (MMRRC) at The Jackson Laboratory, an NIH-funded strain repository, and was donated to the MMRRC by Robert Vassar, Ph.D., Northwestern University. Mice had *ad libitum* access to food and water. Hemizygous 5XFAD mice were used in this study. Genotyping was performed by Transnetyx (Cordova, TN). Animals were sacrificed at 2, 4, and 6 months of age following procedures approved by the BU Institutional Animal Care and Use Committee (IPROTO202300000054). Anesthesia was administered via intraperitoneal injection of a cocktail containing ketamine (100 mg/kg) and xylazine (5 mg/kg). Perfusion was performed using ice-cold phosphate-buffered saline (PBS). Brains were dissected whole and post-fixed in 4% paraformaldehyde in PBS solution at 4°C overnight. After 3 rinses in PBS, brains were cryoprotected in a solution of 30% sucrose in PBS. Brains were embedded in O.C.T. TissueTek (Sakura FineTek USA). Sagittal brain sections were sliced at 16-μm thickness on the cryostat (Epredia, CryoStar NX50). Sections were collected on Superfrost™ slides and stored at -20°C until further use.

### Immunohistochemistry

Brain sections were rehydrated in PBS and stained with ThioS (0.1 mg/mL in 70% ethanol) for 8 minutes at room temperature. Tissues were rinsed twice in 70% ethanol for 3 minutes each, followed by 3 rinses in distilled water, and a 5-minute rinse in PBS. Tissue permeabilization in blocking solution containing PBS, 0.5% Triton X-100, 5% bovine serum albumin (BSA), 5% normal donkey serum, and/or 5% normal goat serum (Sigma-Aldrich, USA) was performed at room temperature for 1-2 hours. Slides were incubated at 4°C for 1-2 nights with the following primary antibodies diluted in blocking solution containing 0.1% Triton X-100: goat anti-IBA-1 (1:200; Novus NB100-1028), rabbit anti-IBA-1 (1:500; Wako 019-19741), mouse anti-6E10 (1:1000; BioLegend 803001), rat anti-Dectin-1 (1:200; Invitrogen MABG-MDECT-2), rabbit anti-CD68 (1:200; Cell Signaling Technology E307V), rat anti-MHC class II (1:750; Sigma-Aldrich MABF33), and Armenian hamster anti-CD11c (1:100 Biorad MCA1369). Tissues were washed 5 times for 5 minutes each in PBS containing 0.1% Triton X-100 before incubation in PBS containing 1:1000 donkey or goat secondary antibodies conjugated with Alexa Fluor 555 (Invitrogen A-31572; Abcam ab150130), Alexa Fluor 594 (Abcam ab150108), and Alexa Fluor 647 (Invitrogen A48272, A31573, or A78967). Slides were rinsed 5 times for 10 minutes each and coverslipped using ProLong™ Gold Antifade Mountant (Thermo Fisher Scientific, USA). Slides were stored overnight at room temperature and protected from light.

### Microscopy and image processing

Immunostained brain sections were imaged on the ZEISS AxioScan 7 slide scanner under 20x objective. Z-stacks were acquired on 4 channels to capture fluorescent signals from ThioS, IBA-1, 6E10, and the phenotypic markers (CD68, Dectin-1, CD11c, and MHCII) excited at 493 nm, 553 nm, 590 nm, and 653 nm, respectively. Emission signals were acquired at 499-527 nm, 583-601 nm, 612-682 nm, and 662-756 nm, respectively. The region of interest (ROI) in the somatosensory cortex and hypothalamus were cropped and processed on Zen 3.5 (ZEISS). Orthogonally projected maximum intensity projections (MIPs) of 7 µm thickness with a width x height of 600 µm x 300 µm was obtained. At least 6 ROIs from each brain region were generated per animal per group.

### Image analysis by CellProfiler

Using CellProfiler (Stirling et al., 2021), we established a pipeline (version 4.2.6) with 5 IdentifyPrimaryObjects modules to segment microglial cells (defined as distinct, non-overlapping cells), microglial clusters (defined as objects without clear boundaries that exceed the detection threshold set for a single microglial cell), diffuse plaques (defined as 6E10^+^ and ThioS^-^ or with faint ThioS^+^ object ≤10µm in diameter) (Shin et al., 2021), dense-core plaques (defined as 6E10^+^ and ThioS^+^ ≥10 µm in diameter) (Bussière et al., 2004), and the microglial phenotypic markers (CD68, Dectin-1, CD11c, and MHCII). For plaque identification, the minimum diameter of the object was set to 30 pixels (= 10 µm). The maximum diameter was unspecified to allow user flexibility in object size detection based on image-specific signals. To distinguish between single and clusters of microglial cells, 2 modules were developed to identify objects labeled with IBA-1. For all modules, the lower and upper threshold limits were determined based on the intensity levels of detected objects in the image. The output data include multiple parameters, such as object counts, area coverage of each object, percentage overlap of markers with IBA-1, and distance between plaques and microglia. Microglia association with diffuse or dense-core plaques was defined by the location of single PAM or PAM clusters within a 50-µm radius from the edge of a segmented plaque. Based on this definition, in plaque-dense cortex, a single PAM or a PAM cluster may be associated with 2 plaques that are less than 100 µm apart. Images and .csv files generated were used for quality control to ensure accurate object segmentation before R codes were executed to compile the .csv files for each experiment for analysis.

### Statistical analysis

For comparison of means between groups, 2-way ANOVA followed by uncorrected Fisher’s Least Significant Difference (LSD) and Tukey’s posthoc tests and data visualization were performed using GraphPad Prism 11 (GraphPad Software, CA, USA). Data are presented as the mean ± standard error of the mean (SEM). Python was used to generate the box plots showing the percentage of CD68 expression in individual microglial cells and perform 2-way ANOVA followed by Fisher’s LSD test and for Mann-Whitney U test of unpaired groups. Statistical significance of *p* < 0.05 are displayed.

## RESULTS

### Female 5XFAD mice exhibited an earlier spike in diffuse and dense-core plaque burden

To determine whether amyloid pathology begins with different kinetics between male and female 5XFAD mice, we analyzed the cortex which is known for rapid plaque deposition and the hypothalamus––a region known for the absence of Aβ plaque––of 2-, 4-, and 6-months-old 5XFAD mice **(Fig. 1A-B)**. As expected, no 6E10 signal was detected in the hypothalamus in all groups analyzed. At 2 months, negligible Aβ plaques were found in the cortex of male 5XFAD, and 11.2-fold higher diffuse plaques were found in female 5XFAD cortex **(Fig. 1C, D, Supplementary Fig. 1)**. At 4 months, the number of diffuse and dense-core plaques was 3.33- and 7.6-fold higher, respectively, in females compared with male 5XFAD mice **(Fig. 1C, D, Supplementary Fig. 1)**. By 6 months, while the burden of diffuse and dense-core plaques in the female cortex appeared to have reached a plateau with no difference to the male cortex, the total plaque count was increased by 1.96-fold in males **(Fig. 1C-E, Supplementary Fig. 1)**. Overall, amyloid pathology progressed slower in the male 5XFAD cortex and showed significance compared to WT only at 6-months-old **(Fig. 1C, D)**.

**Figure 1.**
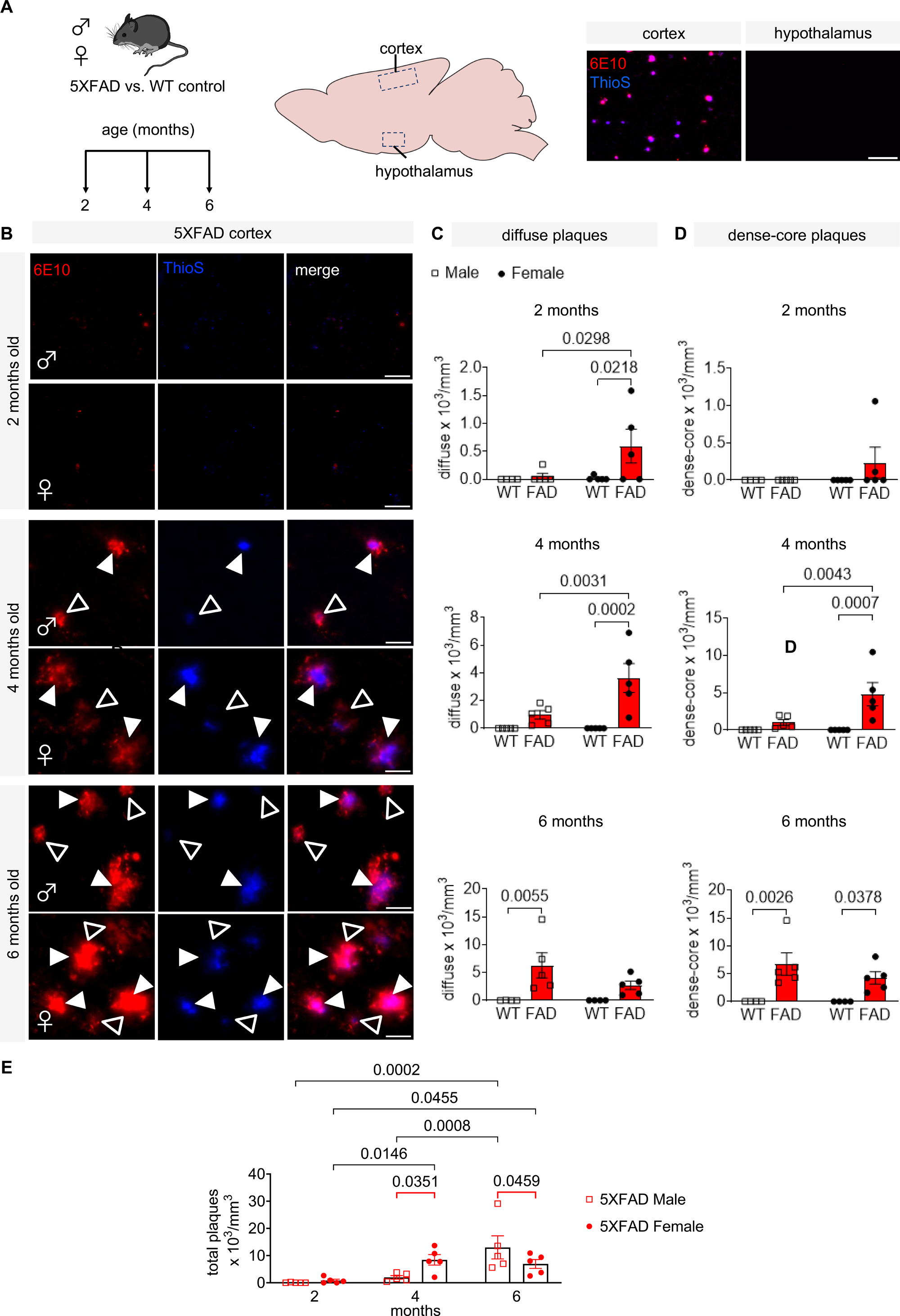
Amyloid-β plaque development in male and female 5XFAD brains. **(A)** Analysis of plaque burden in 5XFAD and WT control cortex and hypothalamus from 2 to 6 months old. Representative overview images of plaque burden in the cortex and hypothalamus in a 4-months-old 5XFAD female mouse. Scale bar, 100 μm. **(B)** Representative images of cortical 6E10^+^ ThioS^−^ diffuse plaques (open arrowheads) and 6E10^+^ ThioS^+^ dense-core plaques (filled arrowheads) in male and female 5XFAD cortex at 2-, 4-, and 6-months-old. 6E10 (red). ThioS (blue). Scale bars, 20 μm. **(C-D)** Quantification of cortical **(C)** diffuse, **(D)** dense-core, and **(E)** total plaques in male (open squares) and female (filled circles) WT (white) and 5XFAD (red). N = 4-5 mice per group. Data are represented as mean ± SEM. Two-way ANOVA and Fisher’s LSD tests between sexes and between genotypes in **(C, D)** or across time in **(E)**. P values < 0.05 are shown.

### Increase in microglial density and cluster formation corresponds with plaque burden, independent of 5XFAD transgenic background

Following the analysis of Aβ plaque load, we wanted to confirm whether microglial proliferation and formation of microglial clusters are specific to PAM. Indeed, no change in the number of single microglial cells was seen in the plaque-free hypothalamus of 5XFAD mice compared to age- and sex-matched WT mice from 2-6 months **(Fig. 2 A, B)**. PAM clusters first appeared at 4 months and at 11.6-fold higher number in the cortex of 4-months-old female 5XFAD mice compared to age-matched 5XFAD males **(Fig. 2 C, D)**. PAM clusters were subsequently observed in the cortex of both sexes by 6 months **(Fig. 2D)**, consistent with progressive increases in 6E10 and ThioS signals **(Fig. 1)**. In addition, 1.47 folds (*p* < 0.0456) more single microglial cells were observed in 6-months-old 5XFAD than WT females **(Fig. 2D)**.

**Figure 2.**
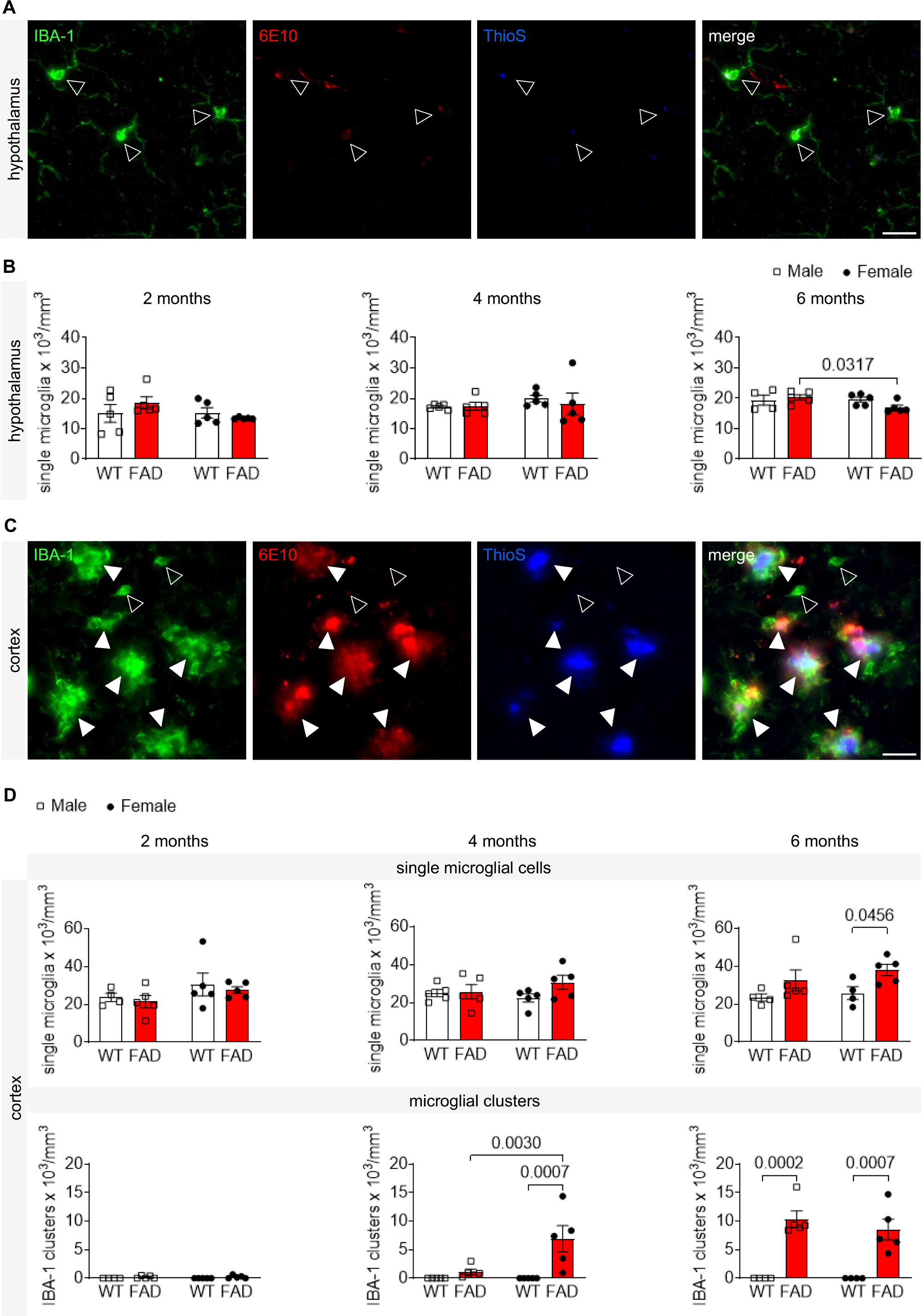
Microglial density and aggregation increase in response to amyloid-β deposition and plaque compaction. **(A)** Representative image of single IBA-1^+^ (green) microglial cells (open arrowheads) and the absence of plaques (6E10, red; ThioS, blue) in the hypothalamus. **(B**) Quantification of single IBA-1^+^ microglial cells in WT and 5XFAD. No microglial cluster was observed in the hypothalamus. **(C)** Representative image of IBA-1^+^ single microglial cells (open arrowheads) and microglial clusters (filled arrowheads) and their proximity to plaques in the cortex. Microglia within 50-µm from the border of a 6E10^+^ plaque are defined as plaque-associated. **(A, C)** Images are from 4-months-old female 5XFAD mice. Scale bars, 20 μm. **(D)** Quantification of cortical IBA-1^+^ single microglial cells and clusters. **(B, D)** Male (squares) and female (circles) WT (white) and 5XFAD (red) brains at 2-, 4-, and 6-months-old. N = 4-5 mice per group. Data are represented as mean ± SEM. Two-way ANOVA and Fisher’s LSD tests between genotypes and between sexes. P values < 0.05 are shown.

### Transient increase in phagolysosomal activity in cortical non-PAM of male 5XFAD before A**β** plaque association

To quantify microglial phagocytic and lysosomal activity (Chistiakov et al., 2017; Jurga et al., 2020), we assessed the levels of CD68, a lysosomal glycoprotein, as a readout for Aβ uptake by non-PAM and PAM 2 to 6 months **(Fig. 3A-C)**. In the absence of Aβ plaque, hypothalamic microglial CD68 expression in 5XFAD and age-matched WT mice maintained a baseline level of 2.98-10.8% in microglia **(Supplementary Fig. 2)**. Interestingly, sex difference in CD68 expression was found in cortical non-PAM in male 5XFAD at 4 months (25.8% in male vs. 8.86% in female 5XFAD, *p* < 0.0430). Phagolysosomal activity was significantly higher in 5XFAD than age-matched 4-months-old WT male microglia (25.8% in 5XFAD vs. 1.45% in WT, *p* < 0.0418) **(Fig. 3B)**. No sex difference in CD68 expression was found in PAM at 4 and 6 months **(Fig. 3C)**.

**Figure 3.**
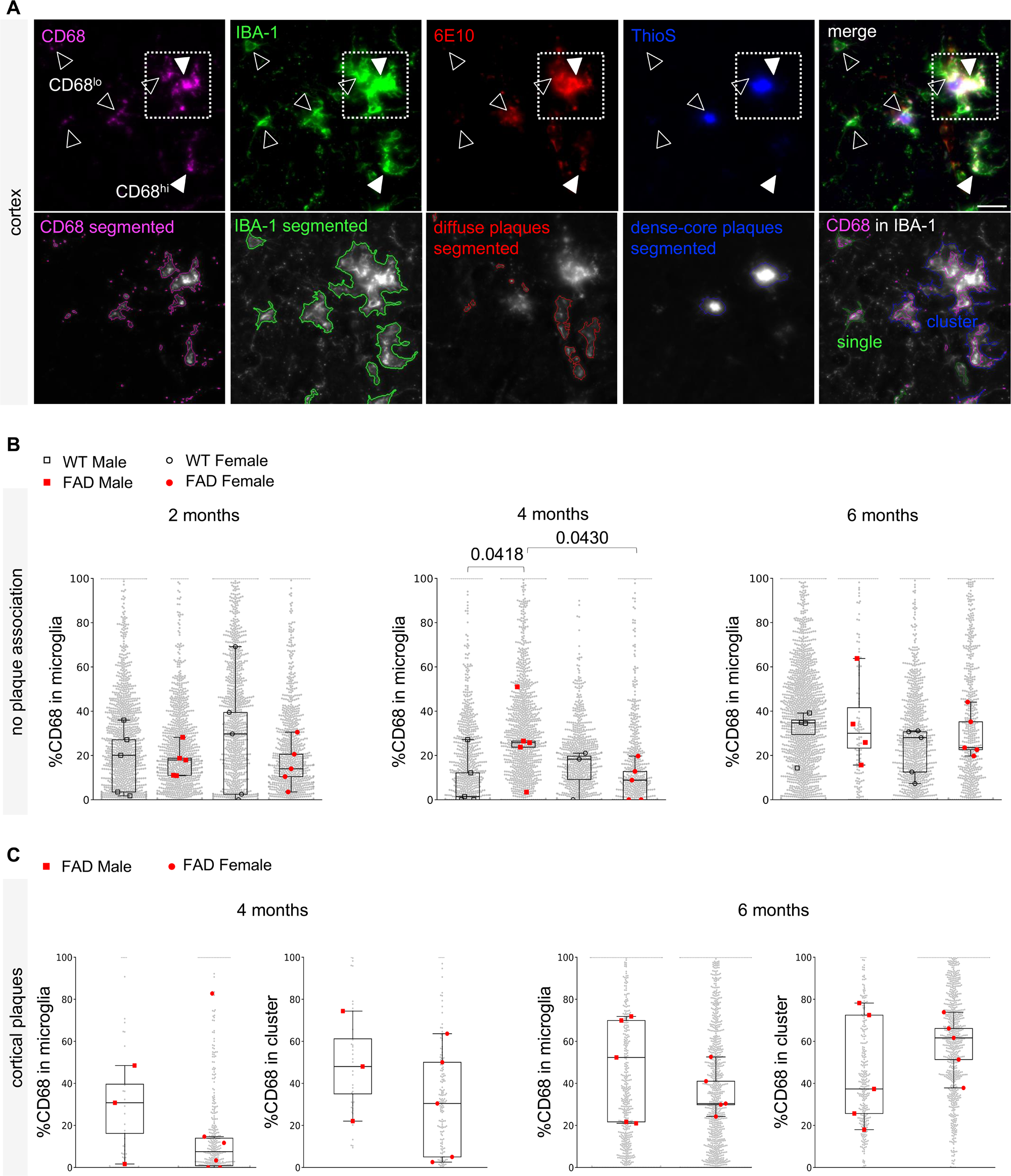
Transient increase in male microglial phagolysosomal activity in 5XFAD cortex before amyloid-β plaque development. **(A)** Representative image of CD68^+^ IBA-1^+^ microglial cells from 4-months-old female 5XFAD cortex. CD68 (magenta; filled arrowheads indicate upregulation in CD68^hi^, open arrowheads indicate low expression in CD68^lo^), IBA-1 (green), 6E10 (red), ThioS (blue), and merge of all upper panels. Boxed region shows a microglial cluster associated with a dense-core plaque that is positive for 6E10 and ThioS. Lower panels show CellProfiler output images indicating the segmentation of each marker and colocalization of CD68 in IBA-1^+^ microglial cells and clusters. Diffuse plaques are 6E10^+^ ThioS^−^ (red outline) and dense-core plaques are 6E10^+^ ThioS^+^ (blue outline). Scale bar, 20 μm. **(B)** Box and swarm plots of percentage CD68 expression (area coverage) in plaque-free single IBA-1^+^ microglial cells (i.e., not associated with plaques) in the cortex of 2-, 4- and 6-months-old WT (black) and 5XFAD (red) mice. N = 3-6 mice represented by 17-24 images per group. **(C)** Box and swarm plots of percentage CD68 expression in plaque-associated cortical IBA-1^+^ single microglial cells and clusters in 4- and 6-months-old 5XFAD (red) mice. N = 5 mice represented by 17-24 images per group. Gray dots indicate the number of cells per group. Squares and circles represent median of percentage CD68 expression per microglial cell of each male and female animal, respectively. Two-way ANOVA and Fisher’s LSD tests between genotypes and between sexes based on the medians. P values < 0.05 are shown.

### Progressive increase of PAM Dectin-1 and CD11c expression during early Aβ pathology

In neurological diseases, microglial Dectin-1/*Clec7a* has been highlighted as a critical factor that mediates Aβ-induced Syk/NFκB signaling, inflammatory cytokine release, and phagocytic synapse-pruning responses (Keren-Shaul et al., 2017; Krasemann et al., 2017; Ulland et al., 2017; Wan et al., 2024; Zhao et al., 2023). Deletion of microglial *Clec7a* in 5XFAD mice led to a twofold increase in plaque burden, exacerbated neuritic dystrophy, and altered the expression of homeostatic neuronal genes (Mangalmurti et al., 2025), suggesting the importance of Dectin-1^+^ microglia in regulating Aβ pathology. Upregulation of CD11c, a cell surface integrin protein commonly used as a marker for activated microglia in developmental apoptosis, neuronal injury and pathology, is associated with phagocytosis (Ghena et al., 2025; Kamphuis et al., 2016; Wlodarczyk et al., 2014; Wlodarczyk et al., 2017a). As amyloid plaque burden rises, microglia localized near plaques have been shown to upregulate both CD11c transcript and protein levels (Kamphuis et al., 2016). Microglia transitioning out of the CD11c-expressing state is understood to suggest a return to homeostatic regulation of lysosomal function and phagocytosis (Ghena et al., 2025).

To further examine the dynamic changes in microglial phagocytic activity during early microglia-plaque interaction, we quantified the levels of microglial Dectin-1 and CD11c in male and female 5XFAD mice at 2-, 4-, and 6-months **(Fig. 4)**. As expected, Dectin-1 is highly upregulated in cortical PAM **(Fig. 4A, B)**. No Dectin-1 or CD11c expression was detected in the hypothalamic microglia of 5XFAD mice and all microglia in WT brains at all time points in the absence of Aβ plaque **(Fig. 4A** and data not shown). From 2–6 months, expression of Dectin-1 in single PAM was gradual in female 5XFAD, whereas marked increases of 32 and 19.4 folds were observed in diffuse and dense-core single PAM, respectively, of male 5XFAD mice **(Fig. 4B)**. During the same period, PAM clusters showed consistent increase in Dectin-1 expression for both sexes by over 58.5 folds and up to 79.4 folds around diffuse and dense-core plaques, respectively **(Fig. 4B)**. The expression of CD11c in PAM was consistently low at 4.48–20.9% in both sexes and showed no sex difference at 6 months **(Fig. 4C, D)**. Our results show no difference in phagocytic activity of PAM between both male and female 5XFAD mice at each time point **(Fig. 4B, D)**. However, the accelerated rate of microglial Dectin-1 expression in male single PAM and PAM clusters over the period of 2–6 months may indicate higher phagocytic and plaque compaction activity leading to the rise in male cortical plaque burden and PAM clusters seen at 6 months **(Figs. 1E, 2D, and 4B)**.

**Figure 4.**
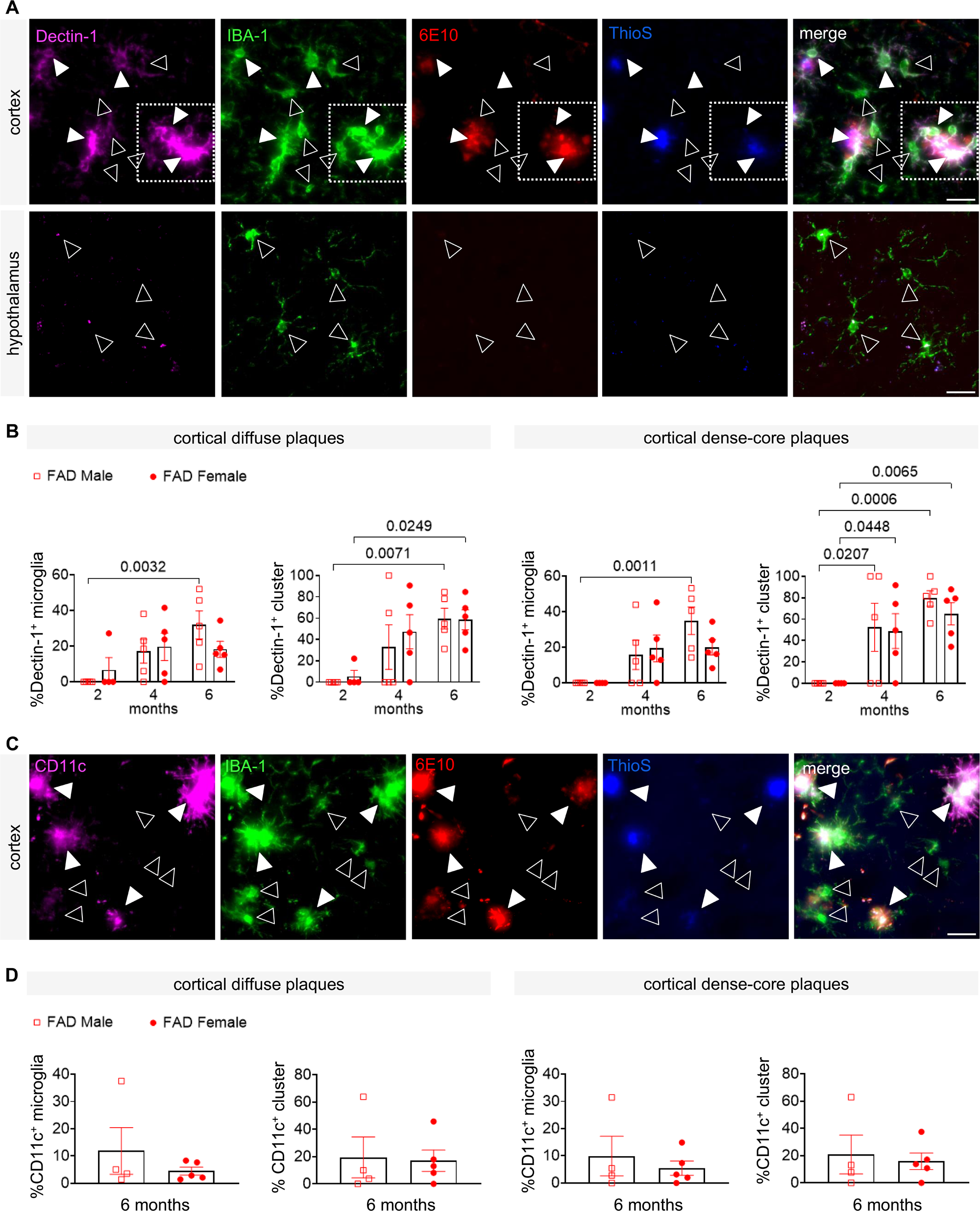
No sex difference in microglial phagocytic activity during early amyloid-β plaque development. **(A)** Representative image of Dectin-1^+^ IBA-1^+^ microglial cells from 4-months-old female 5XFAD cortex. Dectin-1 (magenta; white filled arrowheads for Dectin-1^+^; open arrowheads indicate absence), IBA-1 (green), 6E10 (red), ThioS (blue), and merge of all panels. Boxed region shows a microglial cluster upregulating Dectin-1. No Dectin-1 expression in 5XFAD hypothalamus. Scale bar, 20 μm. **(B)** Percentage of Dectin-1^+^ single microglial cells and microglial clusters associated with cortical diffuse (6E10^+^ ThioS^−^) and dense-core (6E10^+^ ThioS^+^) plaques in 2-, 4-, and 6-months-old 5XFAD male (red squares) and female (red circles) brains. N = 5 mice per group. Data are represented as mean ± SEM. Two-way ANOVA and Tukey’s multiple comparison tests between sex and across time. P values < 0.05 are shown. **(C)** Representative image of CD11c^+^ IBA-1^+^ microglia from 6-months-old female 5XFAD cortex. CD11c (magenta; white filled arrowheads for CD11c^+^; open arrowheads indicate absence), IBA-1 (green), 6E10 (red), ThioS (blue), and merge of all panels. Scale bar, 20 μm. **(D)** Percentage of CD11c^+^ single microglial cells and microglial clusters associated with cortical diffuse (6E10^+^ ThioS^−^) and dense-core (6E10^+^ ThioS^+^) plaques in 6-months-old 5XFAD male (red squares) and female (red circles) brains. No CD11c expression is found in age- and sex-matched WT microglia. N = 5 mice per group. Data are represented as mean ± SEM. Mann Whitney U test.

### MHCII expression remains low in microglia across early plaque stages in both sexes

The loss of MHCII in AD mice was shown to compromise phagocytic capacity and increase amyloid burden (Mittal et al., 2019). Plaque burden was decreased when microglial MHCII in response to Aβ pathology and CD4^+^ T cell signaling was elevated, leading to the conclusion that MHCII performs a neuroprotective role in AD (Mittal et al., 2019). To determine whether antigen-presenting microglia are differentially regulated in male and female 5XFAD mice, we examined the level of MHCII expression in cortical and hypothalamic microglia. As expected, MHCII expression was dependent on the presence of Aβ plaques **(Fig. 5A)**. No MHCII expression was detected in all microglial cells of WT and 5XFAD hypothalamus of both sexes from 2 to 6 months **(Fig. 5A** and data not shown). Our data shows no significant difference between male and female 5XFAD mice in the levels of MHCII expression in single PAM and PAM clusters related to both diffuse and dense-core plaques during the early stages of amyloidosis **(Fig. 5B)**. The number of microglial cells and clusters that upregulate MHCII remained low at 2.44–14.9% by 6 months of amyloidosis and microgliosis. Altogether, this indicates that while MHCII plays a role during AD pathogenesis, early MHCII-mediated PAM response is not integral to the initiation and exacerbation of amyloid pathology early in the disease.

**Figure 5.**
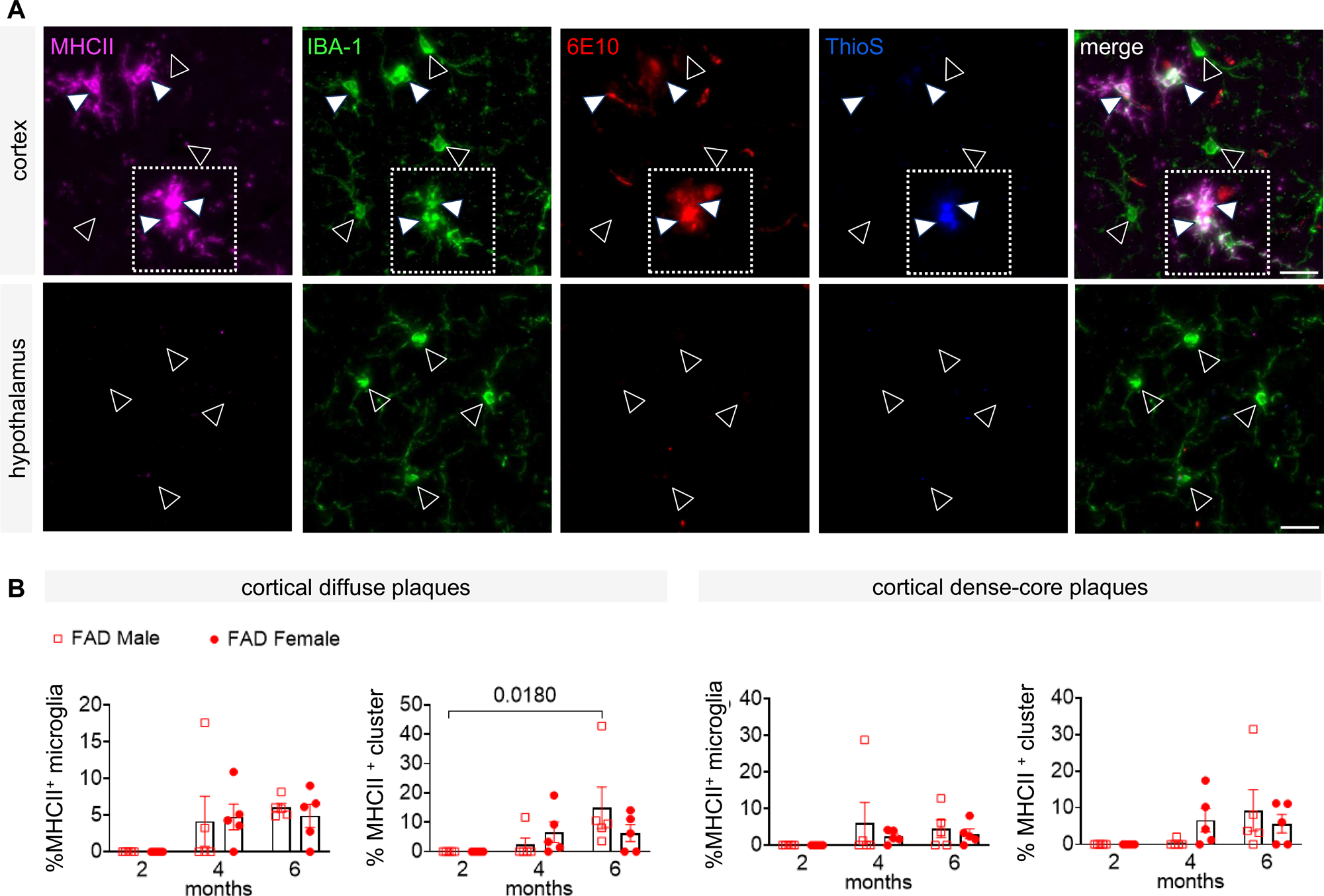
No sex difference in MHC Class II (MHCII) upregulation in early microglial response to amyloid-β plaques. **(A)** Representative images of cortical (top) and hypothalamic (bottom) microglia from 4-months-old female 5XFAD mouse. MHCII (magenta; white filled arrowheads for MHCII^+^; open arrowheads indicate absence), IBA-1 (green), 6E10 (red), ThioS (blue), and merge of all panels. Boxed region shows a microglial cluster upregulating MHCII. No amyloid-β plaques and MHCII expression in microglia are detected in the 5XFAD hypothalamus. Scale bar, 20 μm. **(B)** Quantification of cortical MHCII expression in diffuse and dense-core plaque-associated single microglial cells and clusters from 2-, 4-, and 6-months-old male (red squares) and female (red circles) 5XFAD mice. No MHCII expression is found in age- and sex-matched WT microglia as well as hypothalamic microglia in 5XFAD mice. N = 5 mice per group. Data are represented as mean ± SEM. Two-way ANOVA and Tukey’s multiple comparison tests between sexes and across time. P value < 0.05 is shown.

## DISCUSSION

Our findings demonstrate a pronounced sex difference in microglial response and amyloid load that was transiently observed at 4 months of age during the onset of cortical amyloidosis. Female 5XFAD mice showed a brief spike in diffuse and dense-core plaques that was highly associated with the earlier appearance of microglial clusters compared to male mice. No overt change in microglia residing in the plaque-free hypothalamus of 5XFAD and WT mice were observed between males and females. Our data revealed transient sex difference in non-PAM CD68 levels at 4 months and acceleration of Dectin-1 expression from 2–6 months, further supporting the need to consider sex as a variable in studies of microglial states across brain regions at different stages of health and disease.

### Sex-specific spatiotemporal amyloidogenesis and microgliosis in response to A**β** pathogenesis

We report a direct correlation of a rapid increase in the formation of PAM clusters and cortical dense-core plaques in 4-months-old female 5XFAD mice **(Figs. 1, 2)**. Conversely, slower dense-core plaque formation was found to mirror the delay in the establishment of PAM clusters in male 5XFAD mice of the same age **(Figs. 1, 2)**. Our results do not indicate causality in amyloidogenesis. However, our observation supports the concept that microglia play a significant role in dense-core plaque organization and maturation (Spangenberg et al., 2019). Microglial depletion was shown to reduce the number of dense-core plaques (Lemke and Huang, 2022; Spangenberg et al., 2019). Microglial engulfment of diffuse Aβ deposits, which is dependent on TREM2/DAP12 signaling, has been shown to be essential for compacting plaques to form a protective barrier (Yuan et al., 2016). Microglial aggregation leading to dense-core plaque formation is thought to be neuroprotective as they limit the exposed surface area of neurotoxic Aβ species by sequestering and compacting diffuse amyloid fibrils (Condello et al., 2015; Lemke and Huang, 2022; Yuan et al., 2016). As such, a potentially premature interpretation of our observed plaque and microglial cell density may be that female, rather than male, PAM were more proliferative and thus more efficient in containing Aβ toxicity during disease onset.

### Differences in microglial phagocytic activity correlate with sex-specific progression of amyloid pathology

Our data reveal a sex-dependent, disease stage-specific sequence of microglial activation profiles that may explain the divergent Aβ plaque formation trajectories observed in female and male 5XFAD mice. The higher baseline expression of CD68 in cortical microglia that was observed transiently in male 5XFAD may have enabled effective clearance of nascent Aβ oligomers through heightened phagolysosomal function and resulted in slower plaque formation and expansion before 6 months **(Figs. 1C-E and 3B)**. Testosterone signaling in microglia was shown to promote Aβ-induced autophagy in vitro and in 5XFAD mice, suggesting a role of the sex hormone in mitigating sex-related differences in AD (Du et al., 2025). In contrast, the faster aggregation of Aβ in female 5XFAD cortex appears to promote microgliosis, concomitant PAM upregulation of Dectin-1, and higher plaque compaction than in males before 6 months **(Figs. 1C, 1D, 2D, and 4B)**. While the upregulation of Dectin-1, CD11c, and MHCII is commonly associated with DAM phenotype and progressive neurodegenerative diseases, the physiological and neuroprotective properties of these microglial activation markers have been shown in normal brain development and neurodegenerative or neuroinflammatory diseases where remission occurs (Barclay et al., 2024; Benmamar-Badel et al., 2020; Li et al., 2019; Mittal et al., 2019; Ulland et al., 2017; Wlodarczyk et al., 2014; Wlodarczyk et al., 2017; Wlodarczyk et al., 2019). Taken together, we propose that the increase in PAM expression of these markers at the early stage of amyloidosis could be a protective microglial response mechanism that limits the spread of neurotoxic Aβ throughout the brain. However, without additional supportive interventions, endogenous microglia alone are insufficient to mitigate the systemic impact of multiple AD risk factors. In summary, our findings demonstrating the sex-specific spatiotemporal progression of amyloid pathology and microglial responses emphasize the need to consider both sex and disease stage when developing novel AD therapeutics that target microglial pathways.

## AUTHOR CONTRIBUTIONS

AVM and TLT conceived the study. AVM, AMF, and TLT designed the experiments and performed animal work. AVM, AMF, JT, MI, YM, and ACK processed the samples. AVM and AMF performed microscopy. AMF established the CellProfiler pipelines and wrote the codes with MC. All authors performed data analysis. AVM, AMF, JT, MC, YM, EY, and ACK drafted the manuscript. AVM prepared the figures. TLT supervised the project and edited the manuscript.

## ACKNOWLEDGEMENTS

Drawing of mouse used in Fig. 1 is from the NIH NIAID Visual and Medical Arts library. TLT was supported by a NARSAD Young Investigator Grant from the Brain & Behavior Research Foundation, Spivack Neuroscience Pilot Award, Patricia McLellan Leavitt Research Fund Award, and Boston University startup funding.

## FIGURE LEGENDS

**Supplementary Figure 1.**
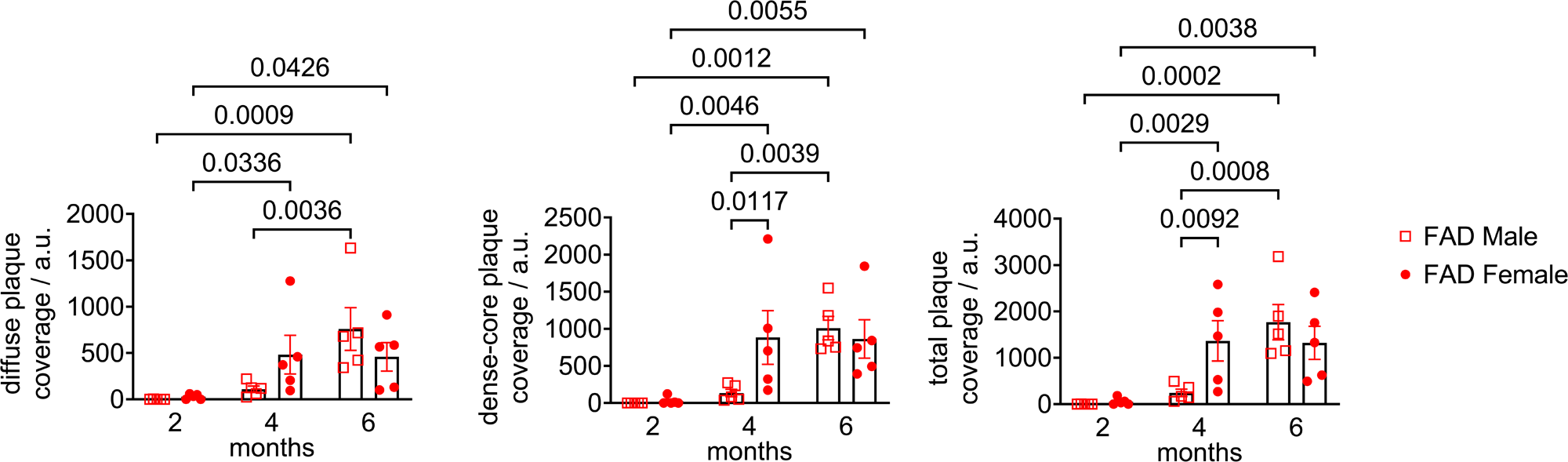
Quantification of amyloid-β plaques in 5XFAD brain cortex. Related to Figure 1. Measurements of diffuse, dense-core, and total plaque area coverage normalized to sampled brain volumes per animal in 2-, 4-, and 6-months-old male (red squares) and female (red circles) mice. N = 5 mice per group. Data are represented as mean ± SEM. Two-way ANOVA and Fisher’s LSD tests between sexes and across time. P values < 0.05 are shown.

**Supplementary Figure 2.**
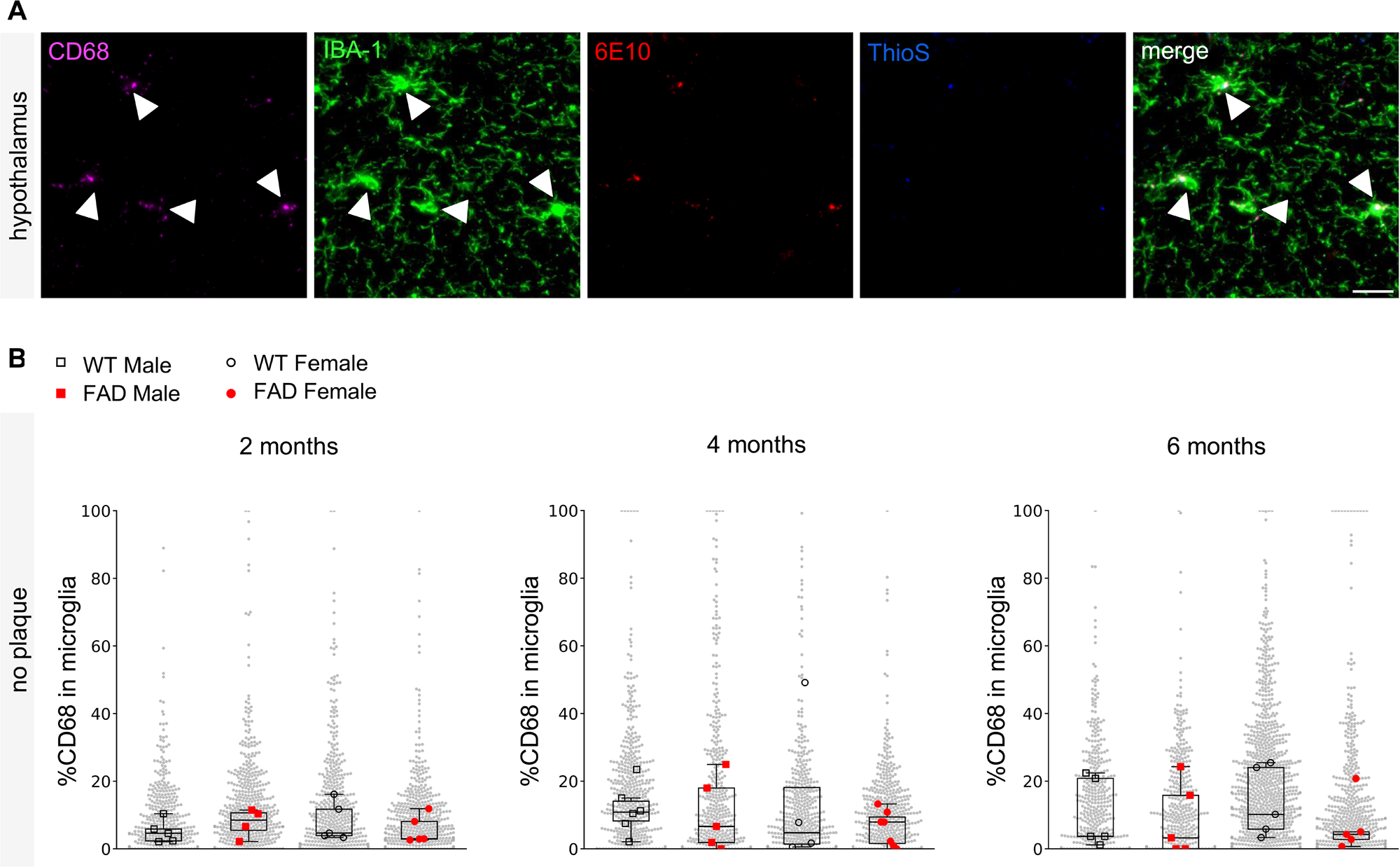
Microglia in the 5XFAD hypothalamus do not upregulate CD68. Related to Figure 3. **(A)** Representative image of CD68^+^ IBA-1^+^ microglial cells from 4-months-old female 5XFAD hypothalamus. No amyloid-β plaques are detected. CD68 (magenta; white arrowheads), IBA-1 (green), 6E10 (red), ThioS (blue), and merge of all upper panels. Scale bar, 20 μm. **(B)** Box and swarm plots of percentage CD68 expression (area coverage) in plaque-free single IBA-1^+^ microglial cells in the hypothalamus of 2-, 4- and 6-months-old WT (black) and 5XFAD (red) mice. N = 4-6 mice represented by 24-27 images per group. Gray dots indicate the number of cells per group. Squares and circles represent median of percentage CD68 expression per microglial cell of each male and female animal, respectively. Two-way ANOVA and Fisher’s LSD tests between genotypes and between sexes based on medians. P values < 0.05 are shown.

